# Stabilization of epithelial β-catenin compromises mammary cell fate acquisition and branching morphogenesis

**DOI:** 10.1101/2023.08.25.554750

**Authors:** Jyoti Prabha Satta, Qiang Lan, Makoto Mark Taketo, Marja Liisa Mikkola

## Abstract

The Wnt/β-catenin pathway plays a critical role in cell fate specification, morphogenesis, and stem cell activation across diverse tissues, including the skin. In mammals, the embryonic surface epithelium gives rise to the epidermis, as well as the associated appendages including hair follicles and mammary glands, both of which depend on epithelial Wnt/β-catenin activity for initiation of their development. Later on, Wnts are thought to enhance mammary gland growth and branching while in hair follicles, they are essential for hair shaft formation. Here we report a strong downregulation of epithelial Wnt/β-catenin activity as the mammary bud progresses to branching. We show that forced activation of epithelial β-catenin severely compromises embryonic mammary gland branching. However, the phenotype of conditional *Lef1* deficient embryos implies that a low level of Wnt/β-catenin activity is necessary for mammary cell survival. Transcriptomic profiling suggests that sustained high β-catenin activity leads to maintenance of mammary bud gene signature at the expense of outgrowth/branching signature. In addition, it leads to upregulation of epidermal differentiation genes. Strikingly, we find a partial switch to hair follicle fate early on upon stabilization of β-catenin suggesting that the level of epithelial Wnt/β-catenin signaling activity may contribute to the choice between skin appendage identities.

## Introduction

The embryonic ectodermal surface epithelium gives rise to the epidermis, as well as ectodermal appendages including hair follicles, sweat and mammary glands, but how the distinct cell fates become allocated in the developing skin remains elusive. In mice, epidermal stratification begins around embryonic day 12.5 (E12.5) and is completed ∼E17.5 when a protective, keratinized barrier has formed (Blanpain and Fuchs, 2009). Concomitant with epidermal morphogenesis, clusters of keratinocytes become fated to skin appendages, each appendage at distinct developmental stage and/or anatomical region (Biggs and Mikkola, 2014; Mikkola and Millar, 2006). In the trunk, mammary glands are the first skin appendages to be induced ∼E11 along the mammary line, a rather narrow ventro-lateral region running between the fore and hind limbs (Spina and Cowin, 2021). Hair follicle anlage are the next ones to appear, the primary ones emerging at E13.5 (Biggs and Mikkola, 2014). As the embryo grows, successive waves of hair follicle induction take place between the existing ones until birth. Sweat gland primordia are the last ones to form, appearing at E17.5 in the ventral paws (Lu and Fuchs, 2014).

Despite the striking differences in the adult organ architecture and function, all ectodermal appendages, including those that develop in the oral cavity such as teeth, share common features that are particularly prominent during the initial stages of development (Biggs and Mikkola, 2014; Mikkola and Millar, 2006). The first morphological sign of a nascent skin appendage is a local epithelial thickening, a placode. Placode formation is accompanied by condensation of the underlying mesenchymal cells. In the next phase, placodes grow into the underlying mesenchyme and transform to spherical buds. Although morphologically similar, the segregation of cellular behaviors becomes notable already at the bud stage: while mammary buds remain largely non-proliferative, the hair buds commence proliferation after a quiescent placode stage (Ahtiainen et al., 2014; Ouspenskaia et al., 2016; Trela et al., 2021).

Following the bud stage, the divergent patterns of epithelial morphogenesis become apparent. In hair follicles, the epithelium grows deeper into the dermis while encapsulating the dermal condensate that matures to dermal papilla, whereas the mammary bud gradually begins to proliferate and sprouts at ∼E15.5 into a secondary mammary mesenchyme, the precursor of the adult fatty stroma, followed by onset of branching morphogenesis a day later (Lan et al., 2023). Mammary rudiments begin to branch as a solid cord of cells, and the bilayered ductal epithelium consisting of inner luminal and outer basal cells characteristic of the adult mammary gland becomes evident only after birth (Hogg et al., 1983; Myllymäki et al., 2023). In hair follicles, differentiation of the multiple concentric cell layers generating the hair shaft and the supporting inner root sheath begins prior to birth, concomitant with the downgrowth of the follicular epithelium (Saxena et al., 2019).

Molecular regulation of skin appendage development has been studied extensively and involves both shared and unique signaling pathways such as the Transforming growth factor (Tgf)β/Bone morphogenetic protein (Bmp), Fibroblast growth factor (Fgf), Ectodysplasin (Eda), Sonic hedgehog (Shh), Parathyroid hormone like hormone (Pthlh, a.k.a. Pthrp), and the Wnt pathway (Lu and Fuchs, 2014; Mikkola and Millar, 2006; Spina and Cowin, 2021). For example, Fgf10/Fgfr2 and Eda signaling pathways are required for the early development of all skin appendages. On the other hand, Shh activity is essential for the growth of the tooth and hair buds, while its suppression is critical for mammary placode formation (Hatsell and Cowin, 2006; Mikkola and Millar, 2006). Pthlh has a unique function in mammary gland development: in its absence, the bud fails to sprout and branch (Wysolmerski et al., 1998).

The canonical Wnt pathway is an evolutionary conserved pathway with essential roles in tissue morphogenesis and cell fate specification in a wide range of organs (Loh et al., 2016). It is mediated by β-catenin, which in unstimulated cells becomes phosphorylated by GSK3β in a ‘destruction complex’ consisting of Axin, Apc, CK1α, and GSK3β priming it for ubiquitination and proteasomal degradation (Nusse and Clevers, 2017; Rim et al., 2022). When a Wnt ligand binds to a receptor complex consisting of a Frizzled family of protein and Lrp5/Lrp6 coreceptor, formation of the destruction complex is inhibited leading to the stabilization and nuclear translocation of β-catenin. In the nucleus, β-catenin interacts with the T cell factor/lymphoid enhancer factor (TCF/LEF) family of transcription factors (Lef1, Tcf7, Tcf7l1, or Tcf7l2, the last three formerly Tcf1, Tcf3, and Tcf4) resulting in the expression of Wnt target genes (Rim et al., 2022).

In developing skin appendages, epithelial Wnt/β-catenin activity is essential for placode formation (Ahn et al., 2013; Andl et al., 2002; Biggs and Mikkola, 2014; Chu et al., 2004; Liu et al., 2008; Zhang et al., 2009). Transgenic overexpression of Dickkopf-1 (Dkk1), a secreted Wnt inhibitor, prevents placode formation altogether, including the mammary anlage (Andl et al., 2002; Chu et al., 2004), whereas deficiency in *Lef1* leads to absence of mammary placodes 2 and 3, and disappearance of the other ones by E15.5 (Boras-Granic et al., 2006; van Genderen et al., 1994). Mammary placodes are smaller also in *Lrp6* deficient mice leading to a smaller sprout that usually fails to branch (Lindvall et al., 2009). On the other hand, mice overexpressing Wnt1 display hyperplasia already at E18.5 leading later on to an overt hyper-branching phenotype and adenocarcinoma (Cunha and Hom, 1996; Tsukamoto et al., 1988). In *ex vivo* cultured embryonic mammary glands, exogenous Wnt3A enhances growth and branching (Voutilainen et al., 2012). All these findings point to a positive role for Wnt/β-catenin pathway in branching morphogenesis. However, the early developmental arrest of mammary primordia in the Dkk1 overexpressing, and *Lef1* null, and conditional β-catenin deficient mouse models has hampered further attempts to uncover the function of the Wnt/β-catenin pathway in embryonic mammary gland development. Moreover, it is often not clear if the epithelium, the mesenchyme, or both are affected, and therefore the precise role and the downstream targets of Wnt/β-catenin signaling in the embryonic mammary epithelium remain largely unknown.

Here, we address these questions by genetically activating β-catenin and conditionally deleting *Lef1* in the embryonic mammary epithelium. Surprisingly, our results point to a negative role for epithelial Wnt/β-catenin signaling in the outgrowth of the mammary anlage, a conclusion supported by the downregulation of all hallmarks of active Wnt/β-catenin signaling in wild type embryos as the bud progressed towards branching. Transcriptomic profiling of the β-catenin activated mammary epithelia revealed suppression of the outgrowth signature at the time of branching despite the initial upregulation of the outgrowth signature at the bud stage. Instead, sustained β-catenin signaling led to ectopic expression of epidermal differentiation genes. Moreover, the mammary bud epithelium adopted a partial hair follicle fate.

## Results

### Size of the mammary bud is tuned by the level of epithelial Wnt/β-catenin activity

To begin to decipher the role of Wnt/β-catenin signaling in embryonic mammary gland (MG) development, we first used a gain-of-function approach and stabilized β-catenin in the epithelium. To this end, we crossed the exon 3 floxed *Ctnnb1* (encoding β-catenin) mice (Harada et al., 1999) with knock-in K14-Cre mice (Huelsken et al., 2001) to generate *Ctnnb1*^Δex3K14/+^ embryos (hereafter stab-β-cat) expressing one wild type and one exon 3 floxed *Ctnnb1* allele lacking the GSK3β phosphorylation sites. The K14-Cre line used is active in the mammary line and mammary buds already at E12.5 (Narhi et al., 2008). As the formation of mammary buds (MB) is asynchronous (Mailleux et al., 2002) we focused our analysis on one anterior (MB2) and one posterior (MB4) mammary primordium and used 3D confocal whole-mount imaging for their analysis. At E12.5, Cre activation manifested as ectopic expression areas of *Ctnnb1* (Supplementary Figure S1a), yet no gross difference was observed between the control and stab-β-cat mammary buds (Figure 1a), a finding in line with the relatively normal expression of bud markers *Wnt10b* and *Edar* at this stage (Narhi et al., 2008). Volume quantification revealed a slight increase in the size of MB2 though (Figure 1b). However, at E13.5, stab-β-cat mammary buds were much larger with precocious sprout-like structures (Figure 1a and b).

**Figure 1.**
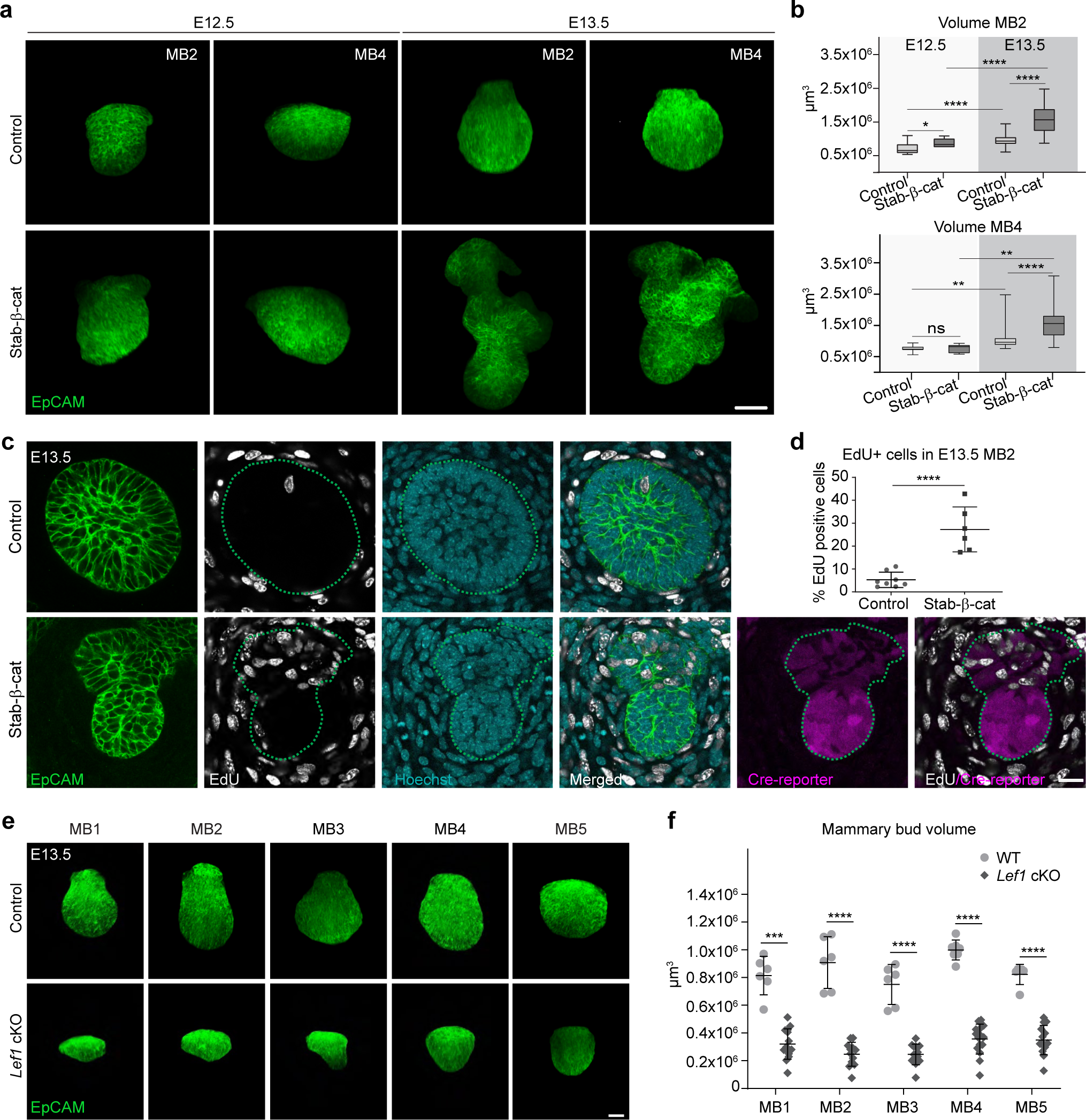
Mammary bud size is tuned by the level of Wnt/β-catenin signaling activity. **(a)** 3D confocal views of E12.5 and E13.5 mammary buds of control and stab-β-cat embryos whole-mount stained with EpCAM. Bar = 30 µm. **(b)** Volume quantification of mammary buds of control and stab-β-cat embryos (E12.5: MB2, n_ctr_ = 17 and n_stab-β-cat_ = 7, and MB4, n_ctr_ = 13 and n_stab-β-cat_ = 6; E13.5: MB2, n_ctr_ = 49 and n_stab-β-cat_ = 46, and MB4, n_ctr_ = 35 and n_stab-β-cat_ = 38). **(c)** Confocal optical sections of MB2 of control and stab-β-cat embryos whole-mount stained with EpCAM, EdU, and Hoechst at E13.5. Dotted line indicates the epithelial-mesenchymal border. Bar = 20 µm. **(d)** Quantification of EdU+ cells in MB2 of control (n = 8) and stab-β-cat (n = 6) embryos at E13.5. **(e)** 3D confocal views of mammary buds of control and *Lef1* cKO embryos whole-mount stained with EpCAM at E13.5. Bar = 30 µm. **(f)** Volume quantification of mammary buds of control and *Lef1* cKO embryos at E13.5 (MB1, n_ctr_ = 6 and n*_Lef1_* _cKO_ = 14; MB2, n_ctr_ = 6 and n*_Lef1_* _cKO_ = 12; MB3, n_ctr_ = 6 and n*_Lef1_* _cKO_ = 13; MB4, n_ctr_ = 7 and n*_Lef1_* _cKO_ = 16; MB5, n_ctr_ = 6 and n*_Lef1_* _cKO_ = 14). Data shown in **(b)** represent the median (line) with 25th and 75th percentiles (hinges) plus the min to max ranges (whiskers). Data in **(d, f)** are shown as mean ± SD. Student’s t-test was used to assess statistical significance. MB, mammary bud; cKO, conditional knockout.

Mammary primordia are relatively non-proliferative until E15.5 (Lan et al., 2023; Trela et al., 2021). Analysis of ethynyl-2′-deoxyuridine (EdU) incorporation at E13.5 showed that in stark contrast to controls, stab-β-cat mammary buds were highly proliferative (Figure 1c and d and Supplementary Figure S1b and c). Analysis of mutant embryos carrying a fluorescent tdTomato (tdT) Cre reporter revealed that the buds often consisted of two distinct regions: those with mosaic and less intense Cre reporter expression and those where most cells expressed high levels of tdT. Intriguingly, the latter were consistently devoid of EdU+ cells, hence resembling control buds (Figure 1c and Supplementary Figure S1b).

Next, we complemented our findings by using a loss-of-function approach by conditionally deleting *Lef1* in the epithelium (hereafter *Lef1* cKO) using a transgenic K14-Cre line (Hafner et al., 2004) ablating Lef1 protein expression by E13.5 (Supplementary Figure S1d). Unlike in the germline deleted *Lef1* embryos (van Genderen et al., 1994), all five pairs of mammary buds formed in *Lef1* cKO embryos (Figure 1e). However, at E13.5 they were all substantially reduced in size compared to control littermates (Figure 1f).

### Sustained β-catenin activation compromises branching

Wnt/β-catenin signaling is commonly regarded as a positive regulator of mammary gland growth and branching (Spina and Cowin, 2021). In wild type embryos, anterior glands begin to branch at E16.5 and the posterior ones by E17.5 (Lindström et al., 2022); Figure 2a and Supplementary Figure S2a). Volume quantification revealed that MG2 grew 3.4-fold in size between E15.5 and E17.5, and by E17.5, on average 5.5 ductal tips had formed in control embryos (Figure 2b and c). At E15.5, MG2 of stab-β-cat embryos appeared as an elongated sprout, but surprisingly, the size was no longer larger than in control embryos (Figure 2a and b). At E16.5 and E17.5, the volume was smaller than in control embryos indicating that growth was stalling and only few branches formed (Figure 2a-c). As analysis of MG4 gave similar results (Supplementary Figure S2a-c), we focused the following quantitative analyses on MG2.

**Figure 2.**
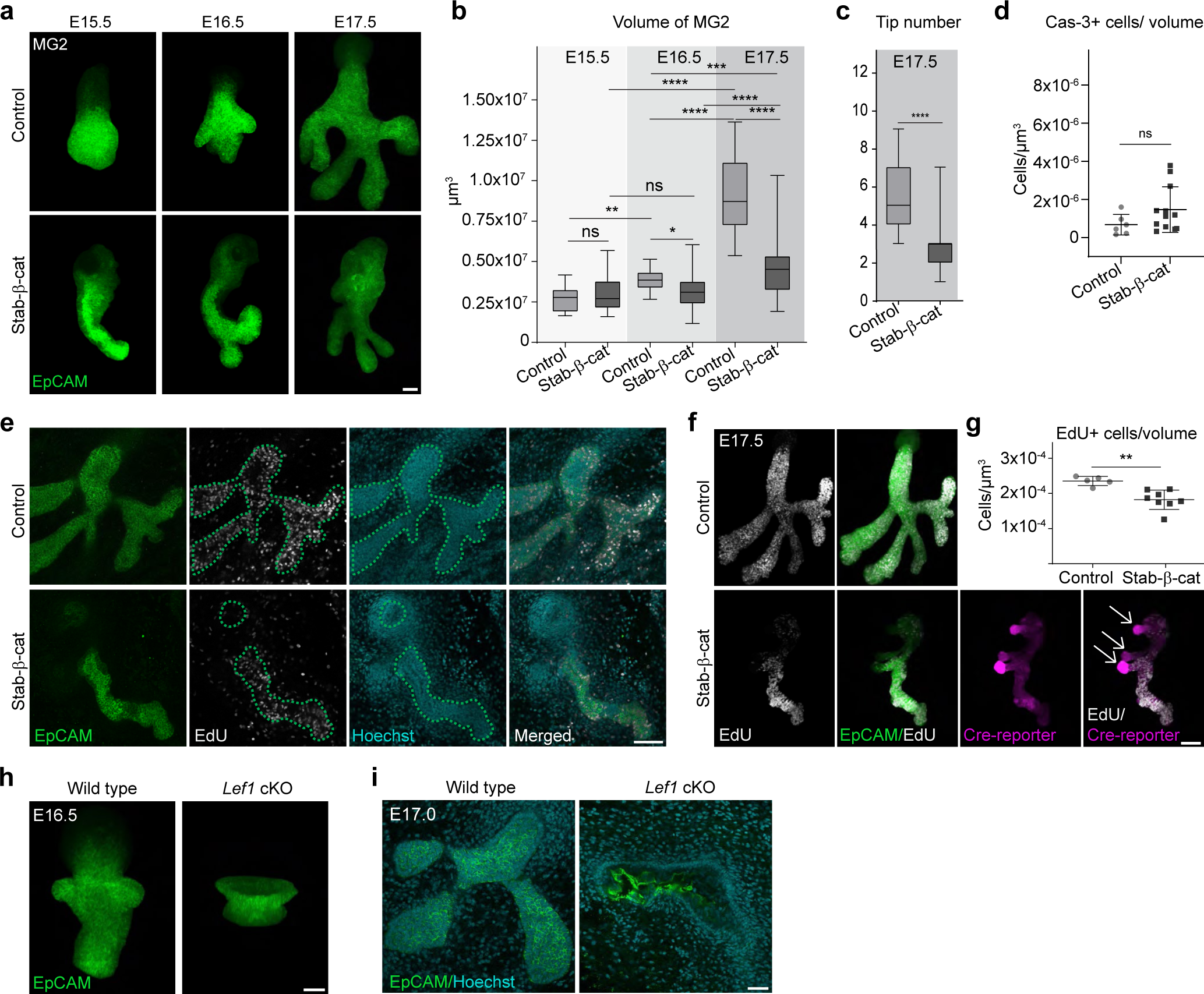
Sustained β-catenin activation compromises growth and branching morphogenesis. **(a)** 3D confocal views of MG2 of E15.5, E16.5, and E17.5 control and stab-β-cat embryos whole-mount stained with EpCAM. Bar = 50 µm. **(b)** Volume quantification of MG2 of control and stab-β-cat embryos (E15.5: n_ctr_ = 14 and n_stab-β-cat_ = 14; E16.5: n_ctr_ = 25 and n_stab-β-cat_ = 30; E17.5: n_ctr_ = 28 and n_stab-β-cat_ = 53). **(c)** Quantification of number of branch tips in MG2 of control (n = 28) and stab-β-cat (n = 53) embryos at E17.5. **(d)** Quantification of Caspase-3+ cells in MG2 of control (n = 6) and stab-β-cat (n = 12) embryos at E17.5. **(e, f)** Confocal optical sections **(e)** and 3D views **(f)** of EdU+ cells in MG2 of control and stab-β-cat embryos at E17.5. Dotted line indicates the epithelial-mesenchymal border in **(e)**, and arrows Cre-reporter positive regions with few EdU+ cells in **(f)**. Bar = 100 µm. **(g)** Quantification of EdU+ cells in MG2 of control (n = 5) and stab-β-cat (n = 8) embryos at E17.5. Bar = 50 µm. **(h)** Representative 3D confocal views of MG1 of control and *Lef1* cKO embryos whole-mount stained with EpCAM at E16.5. **(i)** Representative confocal optical sections of MG2 of control and *Lef1* cKO embryos whole-mount stained with EpCAM and Hoechst at E17.0. In total, six control and four *Lef1* cKO female embryos from four litters were analyzed at E16.5-E17.5. Bar = 50 µm in **(h, i)**. Data shown in **(b, c)** represent the median (line) with 25th and 75th percentiles (hinges) plus the min to max ranges (whiskers). Data in **(d, g)** are shown as mean ± SD. Student’s t-test was used to assess statistical significance. MG, mammary gland; cKO, conditional knockout.

To decipher the cellular mechanism underlying the surprising growth and branching phenotype of stab-β-cat mammary glands, we first assessed apoptosis but found no difference in the number of cleaved-Caspase-3 positive cells between controls and mutants at E17.5 (Figure 2d and Supplementary Figure S2d). Instead, quantification of EdU+ cells revealed a significant reduction in proliferating cells in stab-β-cat mammary glands at E17.5 (Figure 2e-g). Similar to E13.5, we noticed areas where nearly all cells were expressing high levels of the Cre-reporter+ (tdT+) and again these were scarce in EdU+ cells (Figure 2f). In control embryos, the ductal tips were enriched with EdU+ cells, as expected (Figure 2e and f). We also aimed to activate β-catenin only in a subset of cells by using the doxycycline inducible K5-rtTA;TetO-Cre model (Diamond et al., 2000; Perl et al., 2002). Doxycyline was injected at E130-E13.5 and the mammary glands analyzed 4 days later. Control glands branched normally, and Cre-reporter (TdT+) cells were rather evenly distributed along the ductal tree, whereas in mutants, growth and branching were again compromised, and the TdT+ Cre-reporter cells often found as tight clusters (Supplementary Figure S2e).

To investigate how reduced epithelial Wnt/β-catenin activity affects branching, we turned to the *Lef1* cKO mouse model. However, only rudimentary buds (Figure 2h), or disintegrated mammary epithelial tissue (Figure 2i) could be detected at E16.5-E17.0 precluding any further analysis. Thus, similar to *Lef1* germline deficient mice (Boras-Granic et al., 2006; van Genderen et al., 1994), loss of epithelial *Lef1* led to regression of mammary rudiments after the bud stage.

### Epithelial Wnt signaling activity decreases as embryonic mammary gland development progresses to branching

To unveil the molecular events underlying the stab-β-cat phenotype, we took an unbiased approach and profiled the transcriptomes of control and stab-β-cat mammary epithelia at E13.5 and E16.5 by RNA-sequencing (RNA-seq). To minimize the effect of the developmental asynchrony, only anterior mammary buds (MB1-3) were used. Mammary primordia were manually micro-dissected followed by epithelial-mesenchymal separation to obtain the epithelial compartment for RNA-seq. Principal component analysis (PCA) showed that the biological replicates clustered well together (Supplementary Figure S3a). PCA also suggested that the transcriptomes of wild type mammary epithelia undergo large changes between E13.5 and E16.5. However, the E16.5 stab-β-cat samples were most distinct from the other ones (Supplementary Figure S3a). Expectedly, single sample gene set enrichment analysis (ssGSEA) with KEGG database demonstrated that Wnt signaling pathway was significantly upregulated in stab-β-cat mammary epithelia (Figure 3a), also shown by analysis of downstream genes such as *Axin2*, *Lef1*, and *Ctnnb1* (Figure 3b). Intriguingly, the pronounced downregulation of *Axin2* and *Lef1* in control samples from E13.5 to E16.5 (Figure 3b), as well as the GSEA with Hallmark gene sets (Liberzon et al., 2015) (Figure 3c) strongly suggested that Wnt/β-catenin signaling activity is normally downregulated by the time branching begins. Of the Wnt ligands, *Wnt3, Wnt4, Wnt5a, Wnt6, Wnt7a, Wnt7b, Wnt10a,* and *Wnt10b* were expressed at moderate to high levels at E13.5, but all were significantly downregulated at E16.5 (Supplementary Figure S3b). *Tcf* genes other than *Lef1* remained unchanged (Supplementary Figure S3c).

**Figure 3.**
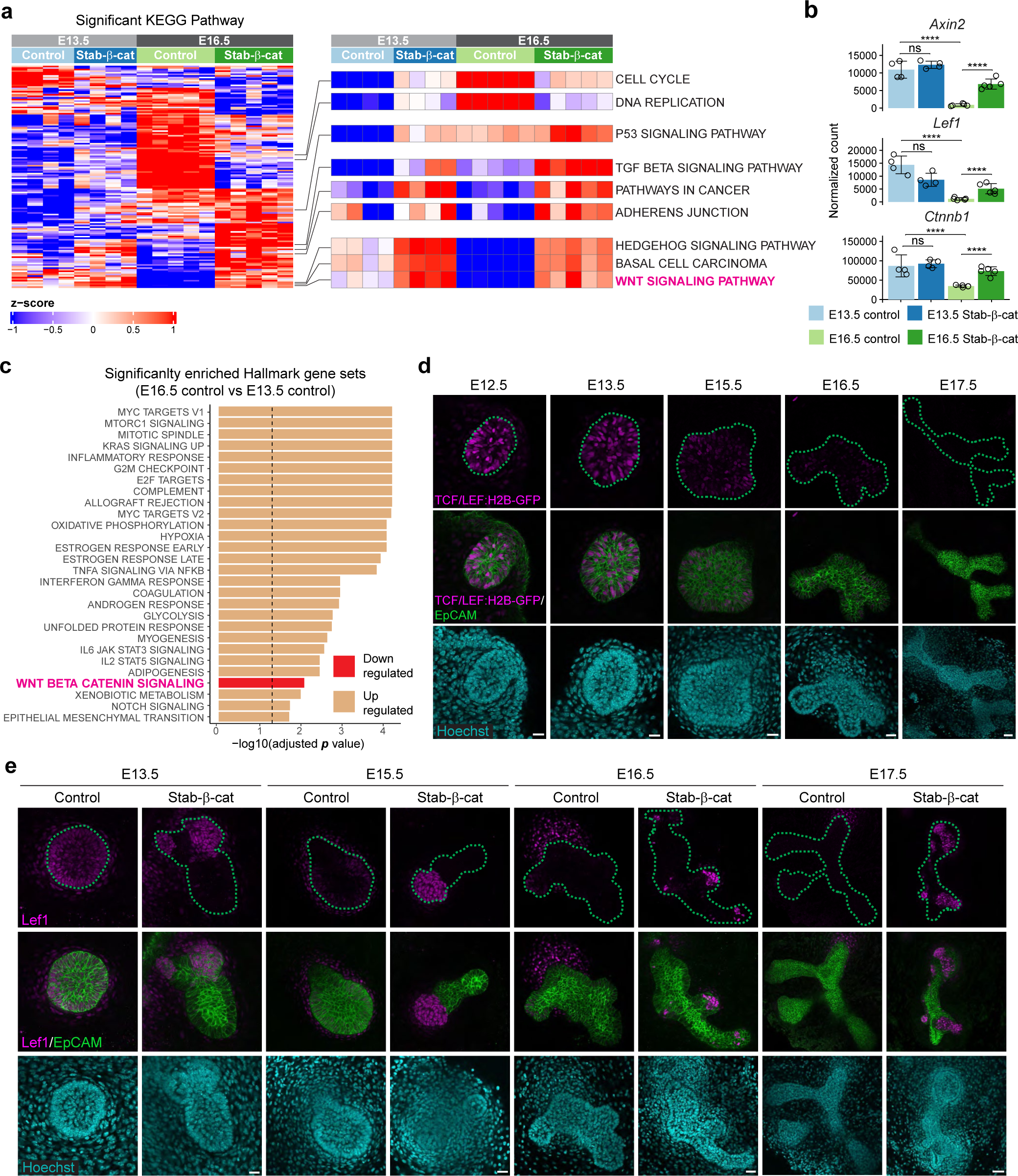
Epithelial Wnt/β-catenin signaling activity decreases as mammary primordia progress to branching. **(a)** Heatmap showing the significantly enriched (adjusted p < 0.01) KEGG pathways comparing E13.5 and E16.5 control and stab-β-cat mammary epithelial transcriptomes by RNA-seq. Selected pathways have been zoomed in. **(b)** Expression of key Wnt pathway genes measured by RNA-seq. Data are shown as mean ± SD. Statistical significance were calculated with Wald test using DESeq2. **(c)** Significantly enriched (adjusted p < 0.05) Hallmark pathways between E13.5 and E16.5 control mammary epithelia. **(d)** Representative confocal optical sections of E12.5-E17.5 mammary primordia (MG2) of transgenic TCF/LEF:H2B-GFP embryos whole-mount stained with EpCAM and Hoechst (n_E12.5_ = 7; n_E13.5_ = 13; n_E15.5_ = 13; n_E16.5_ = 15; n_E17.5_ = 14). Bar = 20 µm (E12.5 and E13.5), 30 µm (E15.5 and E16.5), 50 µm (E17.5). **(e)** Representative confocal optical sections of E13.5-E17.5 control and stab-β-cat mammary primordia whole-mount stained with EpCAM, Hoechst, and Lef1 (E13.5: n_ctr_ = 11 and n_stab-β-cat_ = 5; E15.5: n_ctr_ = 2 and n_stab-β-cat_ = 2; E16.5: n_ctr_ = 5 and n_stab-β-cat_ = 7, and E17.5: n_ctr_ = 9 and n_stab-β-cat_ = 13). Bar = 20 µm (E13.5), 30 µm (E15.5 and E16.5), and 50 µm (E17.5). Dotted line indicates the epithelial-mesenchymal border in **(d, e)**. MB, mammary bud; MG, mammary gland.

To ascertain the downregulation of Wnt/β-catenin signaling activity between the bud and outgrowth stages, we analyzed expression of the transgenic Wnt reporter TCF/LEF:H2B-GFP (Ferrer-Vaquer et al., 2010). Reporter expression was high at the bud stages, but started to decline at E15.5 as the bud progressed to branching, and at E17.5 only few dim H2B-GFP+ cells could be discerned (Figure 3d and Supplemental Figure 3d). Likewise, expression of Lef1, often used as a proxy for Wnt/β-catenin activity, gradually declined as development advanced, and at E17.5 was readily detected only in mesenchymal cells close to the forming nipple (Figure 3e). In contrast, Lef1 expression persisted in stab-β-cat mammary epithelia (Figure 3e) validating the RNA-seq results (Figure 3b). Similar to the Cre-reporter (Figure 2f), cells expressing high levels of Lef1 appeared typically as discrete foci (Figure 3e).

### Stabilization of β-catenin leads to precocious upregulation of mammary outgrowth signature, but to its downregulation when branching begins

To understand the molecular mechanism governing epithelial outgrowth and branching and the impact of stabilized β-catenin therein, we first compared the significantly differentially expressed genes between E13.5 and E16.5 control mammary epithelia. The top-200 genes (excluding lowly expressed genes with average normalized count <100) upregulated at E13.5 were considered as bud signature, and the top-200 genes upregulated at E16.5 as outgrowth (or sprout) signature (Supplementary Table S1). The bud signature included well-known bud markers such as *Dkk4*, *Nrg3*, *Msx1*, and *Wnt10b* (Spina and Cowin, 2021) as well as less-characterized ones like *Sp5*, *Irs4*, *Nos1, Ndnf,* and *Dach2*, the latter two confirmed by *in situ* hybridization (ISH) to be expressed in the mammary buds (Supplementary Figure S4a). The outgrowth signature included genes like *Foxc1, Krt79, Kcnn4, Ly6d, Ets1, Fst, Sox10, Adamts4*, and *Aldh1a3* (Supplementary Table S1). GSEA with Hallmark gene sets revealed that the E16.5 mammary epithelium was positively enriched for genes related to *e.g.* Myc targets, mTORC1 and Kras signaling, as well as cell cycle (Figure 3c).

Next, we analyzed how stabilization of β-catenin affected the wild type bud and outgrowth transcriptomes using gene set enrichment analysis. Strikingly, at E13.5, bud signature genes were significantly downregulated in mutant epithelia (Figure 4a). In total, 83 out of 200 bud signature genes including *Cdh6, Dach2, Epha6, Irs4, Mmp2, Ndnf, Nrg3, and Sema3a* were downregulated (Figure 4b, Supplementary Figure S4b and Supplementary Table S2). Despite the overall downregulation, some bud signature genes (23/200) were upregulated such as *Bmp4*, *Dkk4*, *Fgf20*, *Msx1, Scube1, Prox1, Ascl4, Sp5,* and *Sp6* (Supplementary Figure S4b and c, and Supplementary Table S2), many of them confirmed or suspected transcriptional targets of Wnt/β-catenin (Bazzi et al., 2007; Hu et al., 2013; Narhi et al., 2008; Zhang et al., 2008). Concomitant with the downregulation of the bud signature, we noticed a significant upregulation of the outgrowth signature genes (Figure 4a), in total 84 out of 200 being upregulated (Supplementary Figure S4b), a finding in line with the phenotype analysis showing precocious outgrowth and onset of proliferation in stab-β-cat mammary buds at E13.5 (Figure 1). These genes included e.g., *Aldh1a3, Barx2*, *Ets1, Foxc1, Fst, Kcnn4, Krt79* and *Sox10* (Figure 4c and Supplementary Table S2).

**Figure 4.**
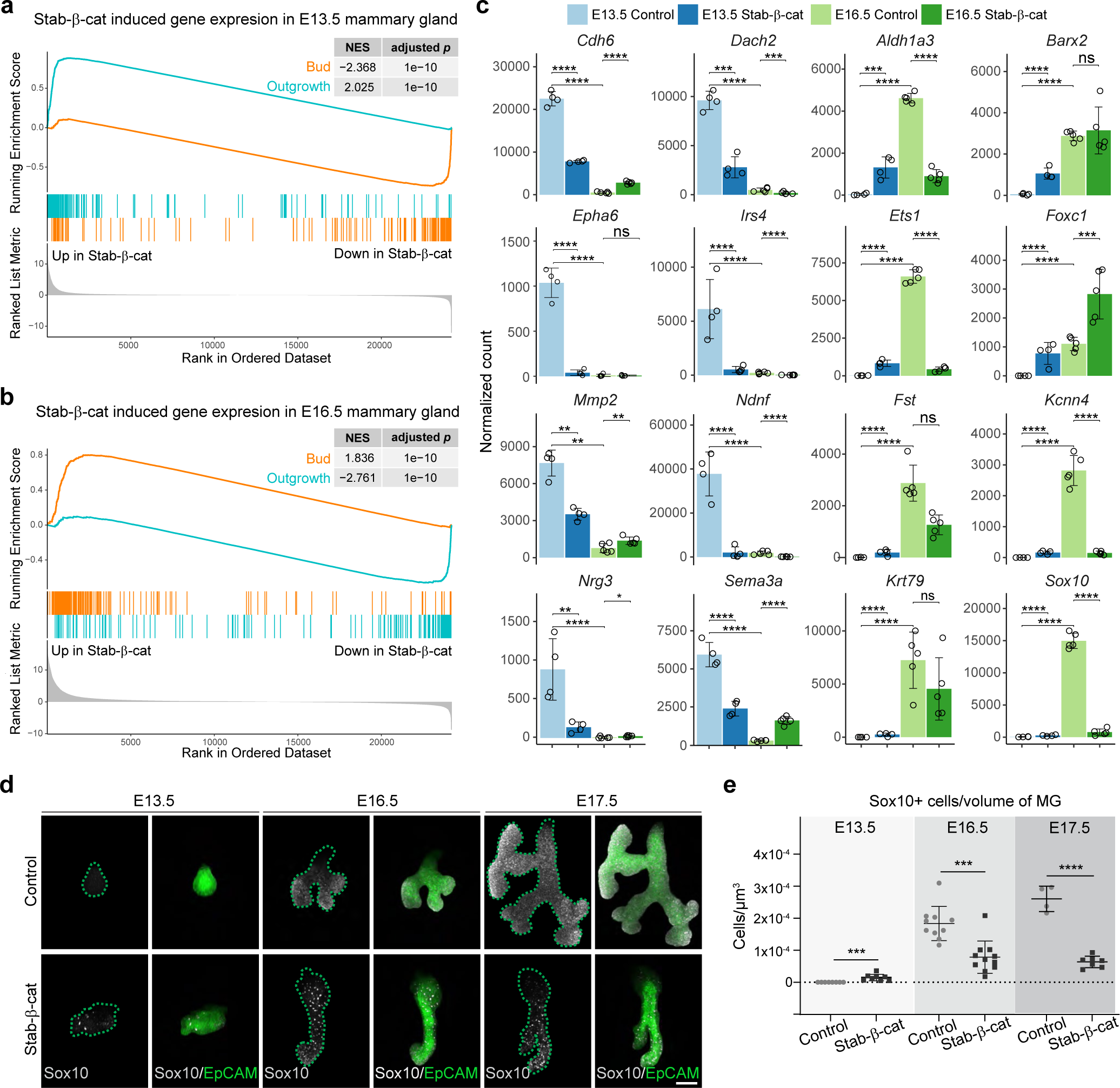
Stabilization of β-catenin leads to precocious upregulation of outgrowth genes, but their downregulation when branching begins. **(a)** Gene set enrichment analysis (GSEA) showing the correlation of the bud and outgrowth transcriptional profiles with genes differentially expressed in stab-β-cat mammary epithelia at E13.5. **(b)** GSEA showing the correlation of the bud and outgrowth transcriptional profiles with genes differentially expressed in stab-β-cat mammary epithelia at E16.5. **(c)** Expression of selected bud and outgrowth signature genes measured by RNA-seq in E13.5 and E16.5 control and stab-β-cat mammary epithelia. Data are shown as mean ± SD. Statistical significance was calculated with Wald test using DESeq2. **(d)** 3D confocal views of E13.5, E16.5, and E17.5 mammary primordia of control and stab-β-cat embryos whole-mount stained with EpCAM and Sox10. Dotted line indicates the epithelial-mesenchymal border. Bar = 100 µm. **(e)** Quantification of Sox10+ cells in control and stab-β-cat mammary primordia (E13.5: n_ctr_ = 11 and n_stab-β-cat_ = 7; E16.5: n_ctr_ = 12 and n_stab-β-cat_ = 13; E17.5: n_ctr_ = 4 and n_stab-β-cat_ = 7). Data are shown as mean ± SD. Student’s t-test was used to assess statistical significance. MG, mammary gland.

In contrast, at E16.5 we observed the exact opposite: upregulation of the bud signature genes (126/200) and downregulation of the outgrowth signature genes (97/200) in stab-β-cat mammary epithelia (Figure 4c and Supplementary Figure S4b). The 126 upregulated bud signature genes included many (40) that were initially downregulated at E13.5 (e.g. *Mmp2, Sema3a*), numerous (63) additional ones such as *Nos1* and *Bdnf*, as well as all the 23 genes that were upregulated already at E13.5 (e.g. *Bmp4*, *Dkk4*, *Fgf20, Msx1, Scube1,* and *Sp5*) (Supplementary Figure S4b and c and Supplementary Table S2) further implying that they may represent direct Wnt/β-catenin target genes. The 97 downregulated outgrowth signature genes included many (33/84) of the same genes that were upregulated at E13.5 such as *Aldh1a3*, *Ets1, Fst, Kcnn4,* and *Sox10*, as well as many additional ones including *Adamts4*, *Lama4*, and *Sfrp1* (Figure 4b, Supplementary Figure 4b and Supplementary Table S1).

For an independent analysis on the effect of β-catenin stabilization on mammary bud enriched genes, we reanalyzed the E12.5 RNA-seq data generated by Kogata *et al*. (Kogata et al., 2018) who had profiled E12.5 mammary bud epithelium and the adjacent non-mammary epidermis allowing us to generate an E12.5 mammary signature of top-200 genes (Supplementary Table S1). Gene set enrichment analysis confirmed downregulation of the E12.5 mammary signature in stab-β-cat epithelia at E13.5, and its upregulation in E16.5 stab-β-cat epithelia (Supplementary Figure S4d and e), in accordance with our own bud signature (Figure 4a and c).

The dramatic downregulation of *Sox10* at E16.5 in stab-β-cat mammary epithelium was particular striking given that deletion of *Sox10* stunts mammary gland outgrowth at ∼E16.5 (Mertelmeyer et al., 2020) and its overexpression drives stemness and invasive behavior (Dravis et al., 2015). To validate our RNA-seq data, we analyzed Sox10 protein expression between E13.5 and E17.5. In wild type embryos, Sox10 was undetectable at E13.5 (Figure 4d and e). At E15.5, a few Sox10+ cells could be detected (Supplementary Figure S4f), and from E16.5 onwards, when branching begins, Sox10 was expressed at high levels (Figure 4d and e). In stab-β-cat embryos, some Sox10+ cells emerged precociously at E13.5 (Figure 4d and e), concordant with the transcriptomic data. Although the number of Sox10+ cells increased substantially thereafter, from E16.5 onwards it remained at much lower levels in mutants compared to control mammary glands (Figure 4d and e).

### Sustained activation of β-catenin upregulates epidermal differentiation program

To gain a more comprehensive understanding on the transcriptional changes downstream of β-catenin stabilization, we performed ssGSEA with Gene Ontology Biological Process (GOBP) terms comparing gene expression of control and stab-β-cat mammary epithelia. Strikingly, the stab-β-cat mammary epithelia were enriched with terms such as skin epidermis development and epidermal cell differentiation (Supplementary Figure S5a and b). At the chromosomal level, many of the significantly differentially expressed genes were located in the same region on chromosome 3 that harbors the epidermal differentiation complex (EDC) locus (Figure 5a and Supplementary Figure 5a). A similar finding was recently made for genes upregulated in Wnt hyperactivated mammary organoids (Mourao et al., 2021). Mourao *et al*. proposed that the squamous differentiation program induced by Wnt/β-catenin could be caused by direct upregulation of epidermal master regulators Grhl3, Klf4, and Ovol1. All these transcription factors were upregulated also in the stab β-cat mutants at E16.5, *Grhl3* and *Klf4* somewhat already at E13.5 (Figure 5b and Supplementary Table S3). Another prominent peak of differentially expressed genes was evident in chromosome 11, enriched with *Krtap* and *Krt* genes, encoding keratin associated proteins and keratins, respectively (Figure 5a and Supplementary Figure 5a). Indeed, analysis of the most differentially expressed genes at E16.5 revealed massive upregulation of EDC locus genes such as *S100a3*, *Sprr1a*, and *Lor* (encoding loricrin), and numerous *Krtap* and *Krt* genes (Figure 5c and Supplementary Table S3). The upregulated *Krt* genes included those involved in epidermal differentiation (e.g. *Krt1*, and *Krt10*), those induced in wounded and stressed epidermis (e.g. *Krt6a*, and *Krt17*), as well as those encoding hard keratins present in the hair shaft and nail (e.g. *Krt31, Krt32*, *Krt81*, and *Krt82*) (Figure 5c and Supplementary Table S3).

**Figure 5.**
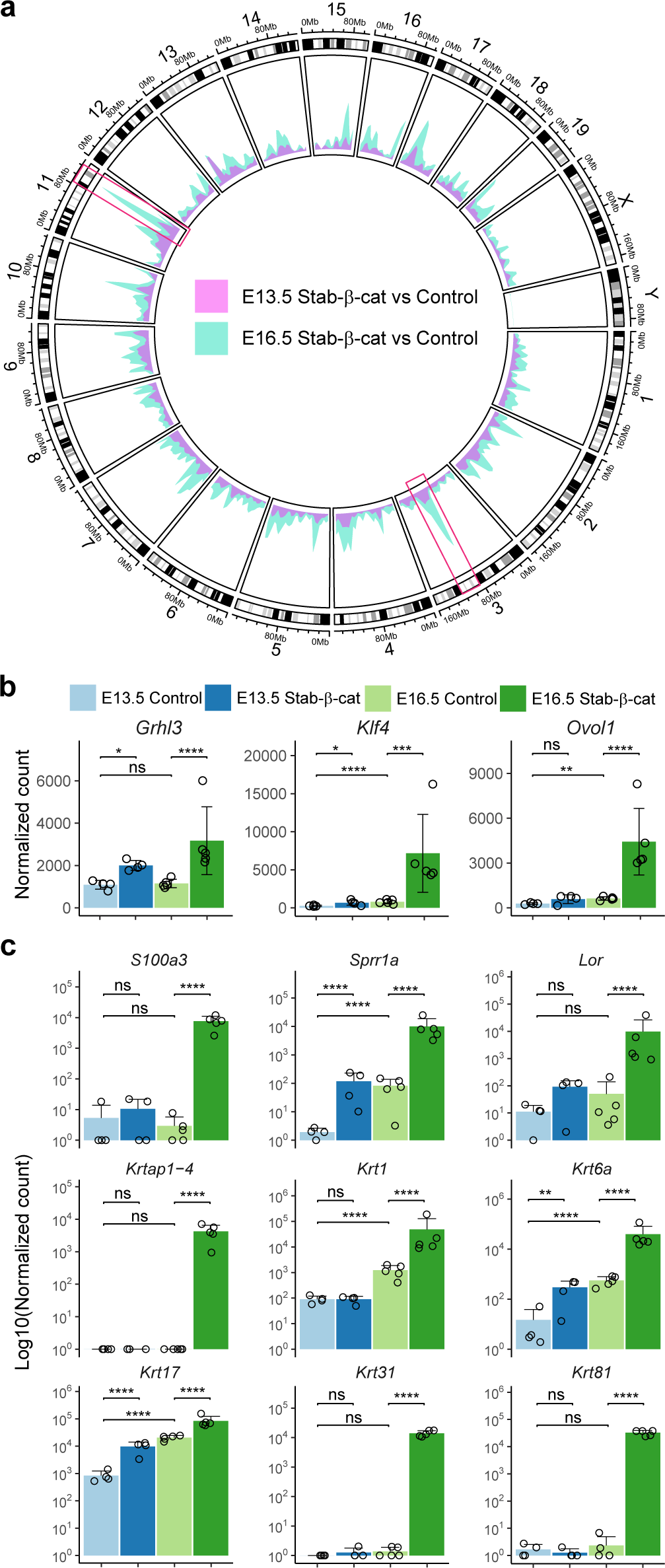
Sustained activation of β-catenin leads to ectopic upregulation of the epidermal differentiation program. **(a)** Circular plot shows the genomic density of DEGs (fold change > 1.5 or < −1.5, adjusted *p* value < 0.05 and average normalized count number > 50) in stab-β-cat versus control at E13.5 or E16.5 mammary gland. Two most striking peaks at chromosomes 3 and 11 are highlighted with a red box. **(b, c)** Expression of selected transcription factors (*Grhl3, Klf4, Ovol1*) **(b)**, and selected genes expressed in the epidermal differentiation complex locus on chromosome 3 (*S100a3, Sprr1a, Lor*), and genes encoding keratin associated proteins or keratins (*Krtap1-4, Krt1, Krt6a, Krt17, Krt31 and Krt81*) **(c)** measured by RNA-seq in control and stab-β-cat mammary epithelia at E13.5 and E16.5. Note Log10 scale used for normalized counts in **(c)**. Data are shown as mean ± SD. Statistical significance was calculated with Wald test using DESeq2.

### Stabilization of β-catenin induces a partial mammary to hair fate switch

In addition to the epidermal differentiation pathways, the ssGSEA of GOBP terms indicated up-regulation of processes such as hair follicle development, regulation of hair cycle, and hair cell differentiation (Supplementary Figure 5a and b), in accordance with upregulation of hard keratin genes. Furthermore, the ssGSEA of KEGG Pathway implicated upregulation of Hedgehog signaling (Figure 3a), normally not active in mammary buds (Hatsell and Cowin, 2006). Accordingly, expression of *Shh*, as well as *Ptch1* and *Gli1*, the commonly used readouts of Hh pathway activity, were significantly upregulated in stab-β-cat mice (Figure 6a). These results prompted us to perform the gene set enrichment analysis of the E13.5 stab-β-cat upregulated genes with our recently identified hair placode enriched transcriptome (Sulic et al., 2023). The analysis confirmed that the hair placode signature (top-200 genes, Supplementary Table S1) was upregulated at E13.5 and E16.5 in stab-β-cat mammary glands (Figure 6b and Supplementary Figure 6c). Comparison with the 142-gene Wnt-high hair bud signature identified by Matos *et al*. (Matos et al., 2020) (Supplementary Table S1) revealed an even more pronounced enrichment in β-catenin mutants already at E13.5, including genes like *Lhx2* (Figure 6c and Supplementary Figure 6d). To validate our findings, we analyzed expression of *Shh* and *Lhx2*, genes normally not expressed in the mammary bud epithelium, *in situ*. Whole mount ISH revealed numerous *Shh*+ foci throughout the mammary line at E13.0 (Figure 6d), and whole mount immunostaining confirmed ectopic expression of Lhx2 in the stab-β-cat mammary bud epithelium (Figure 6e and f).

**Figure 6.**
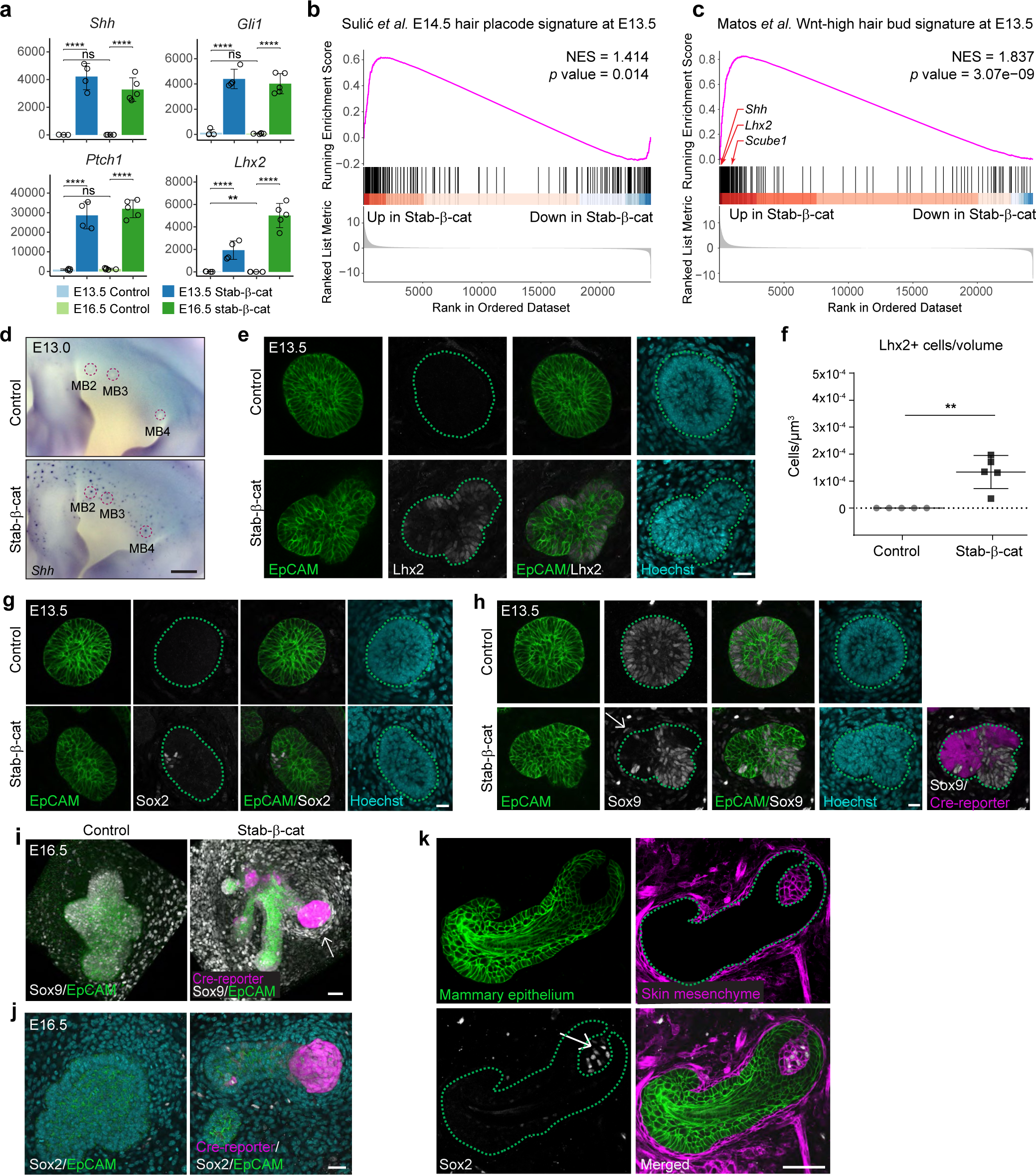
Activation of epithelial β-catenin instigates an early, but partial switch to the hair follicle fate. **(a)** Differential gene expression of *Shh, Gli1, Ptch1* and *Lhx2* measured by RNA-seq. Data are shown as mean ± SD. Statistical significance was calculated with Wald test using DESeq2. **(b)** Gene set enrichment analysis showing correlation between E13.5 control and stab-β-cat with E14.5 hair placode signature defined based on data from Sulić *et al*. (2023). **(c)** Gene set enrichment analysis showing the correlation of genes differentially expressed in the E13.5 stab-β-cat mammary epithelia with the 142-gene Wnt-high hair bud signature from Matos *et al*. (2020). **(d)** Expression of *Shh* was detected by whole-mount *in situ* hybridization in control (n = 8) and stab-β-cat (n = 8) embryos at E13.0. Bar = 500 µm. **(e)** Confocal optical sections of MB2 of control and stab-β-cat embryos whole-mount stained with EpCAM, Lhx2, and Hoechst at E13.5. Bar = 20 µm. **(f)** Quantification of Lhx2+ cells in MB2 of control (n = 5) and stab-β-cat (n = 5) embryos at E13.5. Data are shown as mean ± SD. Student’s t-test was used to assess statistical significance. **(g)** Confocal optical sections of MB2 of control (n = 6) and stab-β-cat (n = 7) embryos whole-mount stained with EpCAM, Sox2, and Hoechst at E13.5. Bar = 20 µm. **(h)** Confocal optical sections of MB2 of control (n = 16) and stab-β-cat (n = 12) embryos whole-mount stained with EpCAM, Sox9, and Hoechst at E13.5. Arrow indicates Sox9+ mesenchymal cells. Bar = 20 µm. **(i)** 3D confocal views of MG2 of control (n = 4) and stab-β-cat (n = 7) embryos whole-mount stained with EpCAM and Sox9 at E16.5. Arrow indicates Sox9+ mesenchymal cells. Bar = 50 µm. **(j)** Confocal optical sections of MG2 of control (n = 6) and stab-β-cat (n = 2) embryos whole-mount stained with EpCAM, Sox2, and Hoechst at E16.5. Bar = 30 µm. **(k)** Representative confocal optical section of an E13.5 mammary bud dissected from an embryo ubiquitously expressing mGFP transplanted on E16.5 skin mesenchyme from an embryo ubiquitously expressing membrane-localized tdTomato (mTmG) and whole-mount stained with Sox2 after 6 days of culture. Arrow indicates Sox2+ dermal condensate cells. Bar = 50 µm. Dotted line indicates the epithelial-mesenchymal border in **(e**, **g**, **h**, **k)**. MB, mammary bud; MG, mammary gland.

These findings suggested that the mammary epithelium might be undergoing a fate switch early on upon β-catenin stabilization. This motivated us to ask whether the mesenchyme of the stab-β-cat mammary buds was also adopting a hair follicle fate and, thus, we analyzed expression of Sox2, a well-known dermal condensate and dermal papilla marker (Driskell et al., 2009). No Sox2 could be detected in control mammary buds (Figure 6g). In the stab-β-cat embryos, we did observe expression of Sox2, but only in sporadic epithelial cells, not in the mesenchyme (Figure 6g). However, we did detect Sox9, another dermal condensate marker (Sennett et al., 2015); hair-gel.net) in the mesenchyme of the mutant, but not control embryos (Figure 6h). Additionally, a patchy loss of Sox9 in the epithelium was evident, colocalizing with Cre-reporter high cells. At E16.5, Sox9 expression was prominent in mesenchymal cells that were adjacent to epithelial Cre-reporter (tdT+) high cell clusters (Figure 6i). These Sox9 positive cells appeared as elongated, tightly packed cells resembling the dermal condensate. However, no Sox2 expression could be detected in the mesenchyme (Figure 6j), indicating a failure to adopt proper dermal condensate fate. These results made us wonder whether mammary epithelial fate might already be fixed at E13.5, precluding full adoption of hair follicle fate leading to a failure in dermal condensate induction. To address this question, we performed tissue recombination experiments between intact E13.5 mammary bud epithelia isolated from embryos ubiquitously expressing tdTomato with E15.5/E16.5 back skin mesenchyme harboring dermal condensates, isolated from embryos ubiquitously expressing mGFP, and cultured the explants for 6 days. Majority of the 40 transplanted mammary buds did not survive or stagnated as small buds, yet some did, apparently if they encountered a dermal condensate. In such cases (7/40 mammary buds), the mammary epithelium transformed into a hair follicle-like appendage encasing a Sox2+ dermal papilla (Figure 6k). These results show that the mammary epithelial fate is still malleable at the bud stage.

## Discussion

The critical role of the Wnt/β-catenin pathway in initiation of mammary gland development has been long recognized, as mice overexpressing Dkk1 lack all signs of mammary placode formation (Chu et al., 2004). In addition, epithelial deletion of β-catenin after placode formation leads to hypoplastic buds (Ahn et al., 2013). We observed a similar phenotype in *Lef1* cKO embryos (where Lef1 was deleted after placode formation) indicating that other Tcf family members cannot substitute for Lef1 in this early epithelial function. However, the function of the epithelial Wnt/β-catenin signaling after the bud stage has remained elusive. In germline deleted *Lef1* null mice, apoptosis is notably increased in the mammary mesenchyme and the hypoplastic buds degenerate and eventually disappear (Boras-Granic et al., 2006; van Genderen et al., 1994), as in *Lef1* cKO embryos. This suggests that Lef1 is essential also for epithelial cell survival. Previous studies have shown that Wnt signaling is critical for adult mammary stem (basal) cell maintenance, at least *in vitro* (Badders et al., 2009; Zeng and Nusse, 2010) suggesting a conserved function across developmental stages.

In contrast, stabilization of β-catenin in the mammary epithelium resulted in precocious outgrowth of the mammary bud albeit with an abnormal shape. At first thought, this finding might indicate that in mammary buds, β-catenin regulates cell proliferation and tissue growth, as it does in many other developmental contexts (Nusse and Clevers, 2017). However, 3D analysis revealed a complex response to Wnt/β-catenin hyperactivation. The mutant mammary buds typically consisted of two types of regions: non-proliferative ones (resembling the wild-type mammary buds) that expressed high levels of the Cre reporter implying that these cells also accumulate high levels of stabilized β-catenin, and highly proliferative regions with fewer and dimmer Cre reporter+ cells. One possible explanation for this observation is that in mutant mammary buds, high levels of β-catenin maintain the low proliferative status of the mammary bud, while simultaneously upregulating the expression of paracrine factors that promote proliferation of the neighboring cells. In support of this hypothesis, in embryos mutant for the negative Wnt pathway regulator Lrp4, epidermal cells between the mammary buds ectopically activate Wnt/β-catenin signaling and arrest proliferation (Ahn et al., 2013). A non-cell autonomous upregulation of proliferation was recently reported in a mouse model where β-catenin was activated in the adult mammary epithelium (Lloyd-Lewis et al., 2022). The reason for the clustering of Cre reporter-high cells is currently unknown, but may be related to sorting of β-catenin hyperactivated cells due to differential cell-cell adhesion, as was proposed to occur in the conditional *Apc* null model of epidermal Wnt/β-catenin overactivation (Matos et al., 2020). Deregulated cell-cell adhesion could also compromise cell motility that is essential for branching morphogenesis (Myllymäki et al., 2023).

The exact function of epithelial Wnt/β-catenin signaling in mammary branching morphogenesis has remained elusive. It is perhaps best understood during pregnancy, where Wnts, in particular Wnt4, regulate side-branching and lobuloalveolar development downstream of progesterone (Brisken et al., 2000; Yu et al., 2016). Accordingly, many mouse mutants with enhanced Wnt/β-catenin activity display a phenotype mimicking early pregnancy state (Bernascone et al., 2019; Bradbury et al., 1995; Imbert et al., 2001; Yu et al., 2016). Epithelial deletion of *Wnt4* leads to a pubertal growth defect, albeit mild, indicating a positive role also during puberty (Rajaram et al., 2015). At this stage, Wnt activity (Axin2-LacZ expression) is enriched in the proliferative bulbous terminal end buds (TEBs), housing the progenitor cells that are the major growth drivers of ductal growth (Van Amerongen et al., 2012). We failed to detect any enrichment of the TCF-Lef:H2B-GFP reporter in the embryonic ductal tips. On the contrary, but in line with analysis of other Wnt reporters (Boras-Granic and Hamel, 2013; Chu et al., 2004; Rajaram et al., 2015), we observed strong downregulation of the signaling activity (TCF-Lef:H2B-GFP reporter, and *Axin2* and *Lef1* expression), when mammary buds progressed to branching. This result suggests that the embryonic function of epithelial Wnt/β-catenin signaling might be fundamentally different from that observed during puberty and pregnancy.

Here we report that sustained activation of β-catenin stagnated epithelial growth from the sprout (E15) stage onward. Based on the phenotype, pathway activity analyses, and the transcriptomic changes revealing suppression of the outgrowth gene signature in stab-β-cat mammary epithelium, we propose that attenuation of Wnt/β-catenin activity after the bud stage is critical for the onset of proliferation and branching. At first look, this conclusion appears contradictory to previous studies indicating a positive role for the canonical Wnt pathway in embryonic mammary gland growth: germline deletion of *Lrp6* stunts growth of the mammary sprout (Lindvall et al., 2009) as does epithelial deletion of *Pygo2*, encoding a Wnt pathway co-activator (Gu et al., 2009). However, as *Lrp6* is also expressed in the mesenchyme and the fat pad is severely affected in *Lrp6* null mice (Lindvall et al., 2009), the contribution of epithelial *Lrp6* in the phenotype is not easy to demarcate, while interpretation of the *Pygo2* mutant phenotype is complicated by the fact that Pygo2 has also β-catenin independent functions (Cantù et al., 2017). Yet, excess of Wnt ligands, either *in vivo* or *ex vivo*, enhances growth (Cunha and Hom, 1996; Voutilainen et al., 2012). However, it is possible that in these models the effect of Wnts is mediated by the mesenchyme. Indeed, we have recently shown that mesenchymal activation of Wnt/β-catenin signaling enhances growth of the mammary epithelium (Lan et al., 2023). Additionally, it is possible that a (very) low level of Wnt activity offers a proliferative advantage to epithelial cells, as was recently shown in mammary organoids (Mourao et al., 2021).

Sustained activation of epithelial β-catenin not only suppressed the mammary outgrowth gene signature, but also activated the epidermal differentiation program. Similar findings have been made also in other mouse models where Wnt/β-catenin has been activated in the mammary epithelium, often resulting in squamous metaplasia (Incassati et al., 2010; Lloyd-Lewis et al., 2022; Miyoshi et al., 2002). In humans, Wnt pathway deregulation is specifically enriched in basal-like tumors, including metaplastic ones with squamous features (Hayes et al., 2008; Yu et al., 2016). Our results suggest that squamous differentiation observed in these mouse models and in Wnt-high metaplastic breast cancer could be a direct consequence of Wnt/β-catenin activation, likely via upregulation of Grhl3, Klf4, and Ovol1 (Mourao et al., 2021). Additionally, Klf4 has been shown to interact directly with β-catenin in non-proliferating suprabasal cells to regulate epidermal differentiation in palmoplantar skin, downstream of Wnt10a (Xu et al., 2017).

Several studies indicate that Wnt/β-catenin activity promotes mammary cell fate acquisition: mice deficient in *Sostdc1* or *Lrp4*, encoding a secreted Wnt pathway inhibitor and its receptor, respectively, have larger mammary buds (Ahn et al., 2013; Närhi et al., 2012) and the Wnt pathway activator lithium chloride induces ectopic mammary bud-like structures *ex vivo* (Chu et al., 2004). Furthermore, formation of supernumerary mammary buds that develop in Eda overexpressing (K14-Eda) mice can be inhibited *ex vivo* with the Wnt pathway inhibitor XAV939 (Voutilainen et al., 2015). However, epithelial Wnt β-catenin activity promotes also hair follicle fate (Gat et al., 1998; Narhi et al., 2008; Song et al., 2018; Zhang et al., 2008) raising the question on how the same pathway can regulate two entirely different cell fates. The induction of mammary and hair placodes differs by timing and location, suggesting that the local signaling environment likely plays an important role in defining the skin appendage identity. Intriguingly, we noticed an early upregulation of hair follicle marker genes in mammary buds upon epithelial stabilization of β-catenin. These included genes like *Shh* and *Lhx2* that are also upregulated in the hair germs of *Apc* null conditional mutants displaying hyperactivation of the Wnt/β-catenin pathway (Matos et al., 2020). We speculate that the level of Wnt/β-catenin signaling activity also plays a role in defining the epithelial cell fate, with higher levels supporting hair follicle identity, in particular the *Shh*-high fate, at the expense of mammary fate. In hair placodes, *Shh* is expressed in a centrally positioned cell population (Sulic et al., 2023). Its activity is essential for specification of the peripherally located Sox9+ hair follicle stem cells, (at least) via its ability to attenuate Wnt signaling, and in turn, high Wnt activity in the Shh+ population inhibits Sox9 expression (Matos et al., 2020; Ouspenskaia et al., 2016; Xu et al., 2015). Our observation that Sox9 expression was suppressed also in the Cre reporter high regions of stab-β-cat mammary buds further supports the conclusion that the mutant mammary buds gained hair follicle-like features. The failure to adopt full hair follicle identity might be attributed to the absence of the hair-specific dermal condensate known to be essential for hair follicle maintenance (Yang and Cotsarelis, 2010). This is supported by our tissue recombination experiment demonstrating the capacity of dermal condensate to convert mammary epithelium into a hair follicle. How the distinct skin appendage fates are specified during embryogenesis remains a fascinating area of future research.

## Materials and Methods

Mouse lines used in this study have been described earlier: *Ctnnb1*^flox3/flox3^ (Harada et al., 1999), K14^Cre/wt^ mice carrying a knock-in of Cre in the *Krt14* locus (Huelsken et al., 2001), transgenic K14-Cre mice (Hafner et al., 2004), Rosa26 tdTomato Cre reporter mice (Jackson labs, Stock 007914), TCF/LEF:H2B-GFP Wnt reporter mice (Jackson labs, stock no. 013752; (Ferrer-Vaquer et al., 2010) mice, bi-transgenic K5-rtTA; TetO-Cre mice (Diamond et al., 2000; Perl et al., 2002), and *Lef1*-floxed mice (Jackson labs, Stock no 000664). The dual fluorescent mGFP;mTmG (R26-mGFP;mTmG) mice were maintained as previously described (Lan et al., 2023). All mouse experiments were approved by local ethics committee and National Animal Experiment Board of Finland. The detailed methods are described in Supplementary Materials and Methods.

## Acknowledgements

We thank Ms. Raija Savolainen for excellent technical assistance, and past and present members of the Mikkola lab, in particular Dr. Vinod Kumar, for advice and fruitful discussions. Mr. Marek Lamos is acknowledged for help with *in situ* hybridization. Mouse studies were carried out with the support of HiLIFE Laboratory Animal Center Core Facility, University of Helsinki. Confocal microscopy was conducted at the Light Microscopy Unit, Institute of Biotechnology, supported by HiLIFE and Biocenter Finland, and RNA sequencing in the DNA Sequencing and Genomics Unit at the Institute of Biotechnology, HiLIFE, University of Helsinki.

## Author Contributions

Conceptualization: JS, MLM; Formal Analysis: JS, QL; Investigation: JS, QL; Methodology: JS, QL, MMT; Project Administration: MLM; Funding acquisition: MLM; Supervision: MLM; Visualization: JS, QL; Writing-Original Draft Preparation: JS, QL, MLM; Writing-Review and Editing: JS, QL, MLM. All authors read and approved the final manuscript.

## References

Ahn Y, Sims C, Logue JM, Weatherbee SD, Krumlauf R. Lrp4 and Wise interplay controls the formation and patterning of mammary and other skin appendage placodes by modulating Wnt signaling. Development 2013;140(3):583–93.

Ahtiainen L, Lefebvre S, Lindfors PH, Renvoisé E, Shirokova V, Vartiainen MK, et al. Directional cell migration, but not proliferation, drives hair placode morphogenesis. Developmental cell 2014;28(5):588–602.

Andl T, Reddy ST, Gaddapara T, Millar SE. WNT signals are required for the initiation of hair follicle development. Developmental cell 2002;2(5):643–53.

Badders NM, Goel S, Clark RJ, Klos KS, Kim S, Bafico A, et al. The Wnt receptor, Lrp5, is expressed by mouse mammary stem cells and is required to maintain the basal lineage. PloS one 2009;4(8):e6594.

Bazzi H, Fantauzzo KA, Richardson GD, Jahoda CA, Christiano AM. The Wnt inhibitor, Dickkopf 4, is induced by canonical Wnt signaling during ectodermal appendage morphogenesis. Developmental biology 2007;305(2):498–507.

Bernascone I, González T, Barea MD, Carabaña C, Hachimi M, Bosch-Fortea M, et al. Sfrp3 modulates stromal–epithelial crosstalk during mammary gland development by regulating Wnt levels. Nature communications 2019;10(1):2481.

Biggs LC, Mikkola ML. Early inductive events in ectodermal appendage morphogenesis. Seminars in cell & developmental biology: Elsevier; 2014. p. 11–21.

Blanpain C, Fuchs E. Epidermal homeostasis: a balancing act of stem cells in the skin. Nature reviews Molecular cell biology 2009;10(3):207–17.

Boras-Granic K, Chang H, Grosschedl R, Hamel PA. Lef1 is required for the transition of Wnt signaling from mesenchymal to epithelial cells in the mouse embryonic mammary gland. Developmental Biology 2006;295(1):219–31.

Boras-Granic K, Hamel PA. Wnt-signalling in the embryonic mammary gland. Journal of mammary gland biology and neoplasia 2013;18:155–63.

Bradbury JM, Edwards PA, Niemeyer CC, Dale TC. Wnt-4 expression induces a pregnancy-like growth pattern in reconstituted mammary glands in virgin mice. Developmental biology 1995;170(2):553–63.

Brisken C, Heineman A, Chavarria T, Elenbaas B, Tan J, Dey SK, et al. Essential function of Wnt-4 in mammary gland development downstream of progesterone signaling. Genes & development 2000;14(6):650–4.

Cantù C, Pagella P, Shajiei TD, Zimmerli D, Valenta T, Hausmann G, et al. A cytoplasmic role of Wnt/β-catenin transcriptional cofactors Bcl9, Bcl9l, and Pygopus in tooth enamel formation. Science signaling 2017;10(465):eaah4598.

Chu EY, Hens J, Andl T, Kairo A, Yamaguchi TP, Brisken C, et al. Canonical WNT signaling promotes mammary placode development and is essential for initiation of mammary gland morphogenesis. 2004.

Cunha GR, Hom YK. Role of mesenchymal-epithelial interactions in mammary gland development. Journal of mammary gland biology and neoplasia 1996;1:21–35.

Diamond I, Owolabi T, Marco M, Lam C, Glick A. Conditional gene expression in the epidermis of transgenic mice using the tetracycline-regulated transactivators tTA and rTA linked to the keratin 5 promoter. Journal of investigative dermatology 2000;115(5):788–94.

Dravis C, Spike BT, Harrell JC, Johns C, Trejo CL, Southard-Smith EM, et al. Sox10 regulates stem/progenitor and mesenchymal cell states in mammary epithelial cells. Cell reports 2015;12(12):2035–48.

Driskell RR, Giangreco A, Jensen KB, Mulder KW, Watt FM. Sox2-positive dermal papilla cells specify hair follicle type in mammalian epidermis. 2009.

Ferrer-Vaquer A, Piliszek A, Tian G, Aho RJ, Dufort D, Hadjantonakis A-K. A sensitive and bright single-cell resolution live imaging reporter of Wnt/ss-catenin signaling in the mouse. BMC developmental biology 2010;10:1–18.

Gat U, DasGupta R, Degenstein L, Fuchs E. De novo hair follicle morphogenesis and hair tumors in mice expressing a truncated β-catenin in skin. Cell 1998;95(5):605–14.

Gu B, Sun P, Yuan Y, Moraes RC, Li A, Teng A, et al. Pygo2 expands mammary progenitor cells by facilitating histone H3 K4 methylation. Journal of Cell Biology 2009;185(5):811–26.

Hafner M, Wenk J, Nenci A, Pasparakis M, Scharffetter-Kochanek K, Smyth N, et al. Keratin 14 Cre transgenic mice authenticate keratin 14 as an oocyte-expressed protein. genesis 2004;38(4):176–81.

Harada N, Tamai Y, Ishikawa T-o, Sauer B, Takaku K, Oshima M, et al. Intestinal polyposis in mice with a dominant stable mutation of the β-catenin gene. The EMBO journal 1999;18(21):5931–42.

Hatsell SJ, Cowin P. Gli3-mediated repression of Hedgehog targets is required for normal mammary development. 2006.

Hayes MJ, Thomas D, Emmons A, Giordano TJ, Kleer CG. Genetic changes of Wnt pathway genes are common events in metaplastic carcinomas of the breast. Clinical Cancer Research 2008;14(13):4038–44.

Hogg N, Harrison C, Tickle C. Lumen formation in the developing mouse mammary gland. Development 1983;73(1):39–57.

Hu W, Ye Y, Zhang W, Wang J, Chen A, Guo F. miR-142-3p promotes osteoblast differentiation by modulating Wnt signaling. Molecular Medicine Reports 2013;7(2):689–93.

Huelsken J, Vogel R, Erdmann B, Cotsarelis G, Birchmeier W. β-Catenin controls hair follicle morphogenesis and stem cell differentiation in the skin. Cell 2001;105(4):533–45.

Imbert A, Eelkema R, Jordan S, Feiner H, Cowin P. ΔN89β-catenin induces precocious development, differentiation, and neoplasia in mammary gland. The Journal of cell biology 2001;153(3):555–68.

Incassati A, Chandramouli A, Eelkema R, Cowin P. Key signaling nodes in mammary gland development and cancer: β-catenin. Breast Cancer Research 2010;12:1–14.

Kogata N, Bland P, Tsang M, Oliemuller E, Lowe A, Howard BA. Sox9 regulates cell state and activity of embryonic mouse mammary progenitor cells. Communications biology 2018;1(1):228.

Lan Q, Trela E, Lindstrom R, Satta J, Christensen MM, Holzenberger M, et al. On growth and form of the mammary gland: Epithelial-mesenchymal interactions in embryonic mammary gland development. bioRxiv 2023:2023.04. 24.538064.

Liberzon A, Birger C, Thorvaldsdóttir H, Ghandi M, Mesirov JP, Tamayo P. The molecular signatures database hallmark gene set collection. Cell systems 2015;1(6):417–25.

Lindström R, Satta JP, Myllymäki S-M, Lan Q, Trela E, Prunskaite-Hyyryläinen R, et al. Unraveling the principles of mammary gland branching morphogenesis. bioRxiv 2022:2022.08. 23.504958.

Lindvall C, Zylstra CR, Evans N, West RA, Dykema K, Furge KA, et al. The Wnt co-receptor Lrp6 is required for normal mouse mammary gland development. PloS one 2009;4(6):e5813.

Liu F, Chu EY, Watt B, Zhang Y, Gallant NM, Andl T, et al. Wnt/β-catenin signaling directs multiple stages of tooth morphogenesis. Developmental biology 2008;313(1):210–24.

Lloyd-Lewis B, Gobbo F, Perkins M, Jacquemin G, Huyghe M, Faraldo MM, et al. In vivo imaging of mammary epithelial cell dynamics in response to lineage-biased Wnt/β-catenin activation. Cell Reports 2022;38(10):110461.

Loh KM, van Amerongen R, Nusse R. Generating cellular diversity and spatial form: Wnt signaling and the evolution of multicellular animals. Developmental cell 2016;38(6):643–55.

Lu C, Fuchs E. Sweat gland progenitors in development, homeostasis, and wound repair. Cold Spring Harbor perspectives in medicine 2014;4(2):a015222.

Mailleux AA, Spencer-Dene B, Dillon C, Ndiaye D, Savona-Baron C, Itoh N, et al. Role of FGF10/FGFR2b signaling during mammary gland development in the mouse embryo. 2002.

Matos I, Asare A, Levorse J, Ouspenskaia T, de la Cruz-Racelis J, Schuhmacher L-N, et al. Progenitors oppositely polarize WNT activators and inhibitors to orchestrate tissue development. Elife 2020;9:e54304.

Mertelmeyer S, Weider M, Baroti T, Reiprich S, Fröb F, Stolt CC, et al. The transcription factor Sox10 is an essential determinant of branching morphogenesis and involution in the mouse mammary gland. Scientific reports 2020;10(1):17807.

Mikkola ML, Millar SE. The mammary bud as a skin appendage: unique and shared aspects of development. Journal of mammary gland biology and neoplasia 2006;11:187–203.

Miyoshi K, Shillingford JM, Le Provost F, Gounari F, Bronson R, Von Boehmer H, et al. Activation of β-catenin signaling in differentiated mammary secretory cells induces transdifferentiation into epidermis and squamous metaplasias. Proceedings of the National Academy of Sciences 2002;99(1):219–24.

Mourao L, Zeeman AL, Wiese KE, Bongaarts A, Oudejans LL, Martinez IM, et al. Hyperactive WNT/CTNNB1 signaling induces a competing cell proliferation and epidermal differentiation response in the mouse mammary epithelium. bioRxiv 2021:2021.06. 22.449461.

Myllymäki S-M, Kaczyńska B, Lan Q, Mikkola ML. Spatially coordinated cell cycle activity and motility govern bifurcation of mammary branches. Journal of Cell Biology 2023;222(9).

Narhi K, Jarvinen E, Birchmeier W, Taketo MM, Mikkola ML, Thesleff I. Sustained epithelial β-catenin activity induces precocious hair development but disrupts hair follicle down-growth and hair shaft formation. 2008.

Nusse R, Clevers H. Wnt/β-catenin signaling, disease, and emerging therapeutic modalities. Cell 2017;169(6):985–99.

Närhi K, Tummers M, Ahtiainen L, Itoh N, Thesleff I, Mikkola ML. Sostdc1 defines the size and number of skin appendage placodes. Developmental biology 2012;364(2):149–61.

Ouspenskaia T, Matos I, Mertz AF, Fiore VF, Fuchs E. WNT-SHH antagonism specifies and expands stem cells prior to niche formation. Cell 2016;164(1):156–69.

Perl A-KT, Wert SE, Nagy A, Lobe CG, Whitsett JA. Early restriction of peripheral and proximal cell lineages during formation of the lung. Proceedings of the National Academy of Sciences 2002;99(16):10482–7.

Rajaram RD, Buric D, Caikovski M, Ayyanan A, Rougemont J, Shan J, et al. Progesterone and W nt4 control mammary stem cells via myoepithelial crosstalk. The EMBO journal 2015;34(5):641–52.

Rim EY, Clevers H, Nusse R. The Wnt pathway: from signaling mechanisms to synthetic modulators. Annual review of biochemistry 2022;91:571–98.

Saxena N, Mok KW, Rendl M. An updated classification of hair follicle morphogenesis. Experimental dermatology 2019;28(4):332–44.

Sennett R, Wang Z, Rezza A, Grisanti L, Roitershtein N, Sicchio C, et al. An integrated transcriptome atlas of embryonic hair follicle progenitors, their niche, and the developing skin. Developmental cell 2015;34(5):577–91.

Song Y, Boncompagni AC, Kim S-S, Gochnauer HR, Zhang Y, Loots GG, et al. Regional control of hairless versus hair-bearing skin by Dkk2. Cell reports 2018;25(11):2981–91. e3.

Spina E, Cowin P. Embryonic mammary gland development. Seminars in cell & developmental biology: Elsevier; 2021. p. 83–92.

Sulic A-M, Roy RD, Papagno V, Lan Q, Saikkonen R, Jernvall J, et al. Transcriptomic landscape of early hair follicle and epidermal development. Cell Reports 2023;42(6):112643.

Trela E, Lan Q, Myllymaki SM, Villeneuve C, Lindstrom R, Kumar V, et al. Cell influx and contractile actomyosin force drive mammary bud growth and invagination. J Cell Biol 2021;220(8).

Tsukamoto AS, Grosschedl R, Guzman RC, Parslow T, Varmus HE. Expression of the int-1 gene in transgenic mice is associated with mammary gland hyperplasia and adenocarcinomas in male and female mice. Cell 1988;55(4):619–25.

Van Amerongen R, Bowman AN, Nusse R. Developmental stage and time dictate the fate of Wnt/β-catenin-responsive stem cells in the mammary gland. Cell stem cell 2012;11(3):387–400.

van Genderen C, Okamura RM, Farinas I, Quo R-G, Parslow TG, Bruhn L, et al. Development of several organs that require inductive epithelial-mesenchymal interactions is impaired in LEF-1-deficient mice. Genes & development 1994;8(22):2691–703.

Voutilainen M, Lindfors PH, Lefebvre S, Ahtiainen L, Fliniaux I, Rysti E, et al. Ectodysplasin regulates hormone-independent mammary ductal morphogenesis via NF-κB. Proceedings of the National Academy of Sciences 2012;109(15):5744–9.

Voutilainen M, Lindfors PH, Trela E, Lönnblad D, Shirokova V, Elo T, et al. Ectodysplasin/NF-κB promotes mammary cell fate via Wnt/β-catenin pathway. PLoS genetics 2015;11(11):e1005676.

Wysolmerski JJ, Philbrick WM, Dunbar ME, Lanske B, Kronenberg H, Karaplis A, et al. Rescue of the parathyroid hormone-related protein knockout mouse demonstrates that parathyroid hormone-related protein is essential for mammary gland development. Development 1998;125(7):1285–94.

Xu M, Horrell J, Snitow M, Cui J, Gochnauer H, Syrett CM, et al. WNT10A mutation causes ectodermal dysplasia by impairing progenitor cell proliferation and KLF4-mediated differentiation. Nature communications 2017;8(1):15397.

Xu Z, Wang W, Jiang K, Yu Z, Huang H, Wang F, et al. Embryonic attenuated Wnt/β-catenin signaling defines niche location and long-term stem cell fate in hair follicle. elife 2015;4:e10567.

Yang C-C, Cotsarelis G. Review of hair follicle dermal cells. Journal of dermatological science 2010;57(1):2–11.

Yu QC, Verheyen EM, Zeng YA. Mammary development and breast cancer: a Wnt perspective. Cancers 2016;8(7):65.

Zeng YA, Nusse R. Wnt proteins are self-renewal factors for mammary stem cells and promote their long-term expansion in culture. Cell stem cell 2010;6(6):568–77.

Zhang Y, Andl T, Yang SH, Teta M, Liu F, Seykora JT, et al. Activation of β-catenin signaling programs embryonic epidermis to hair follicle fate. 2008.

Zhang Y, Tomann P, Andl T, Gallant NM, Huelsken J, Jerchow B, et al. Reciprocal requirements for EDA/EDAR/NF-κB and Wnt/β-catenin signaling pathways in hair follicle induction. Developmental cell 2009;17(1):49–61.

